## Supplemental Figures for "Temporal dynamics of nitrogen cycle gene diversity in a hyporheic microbiome"

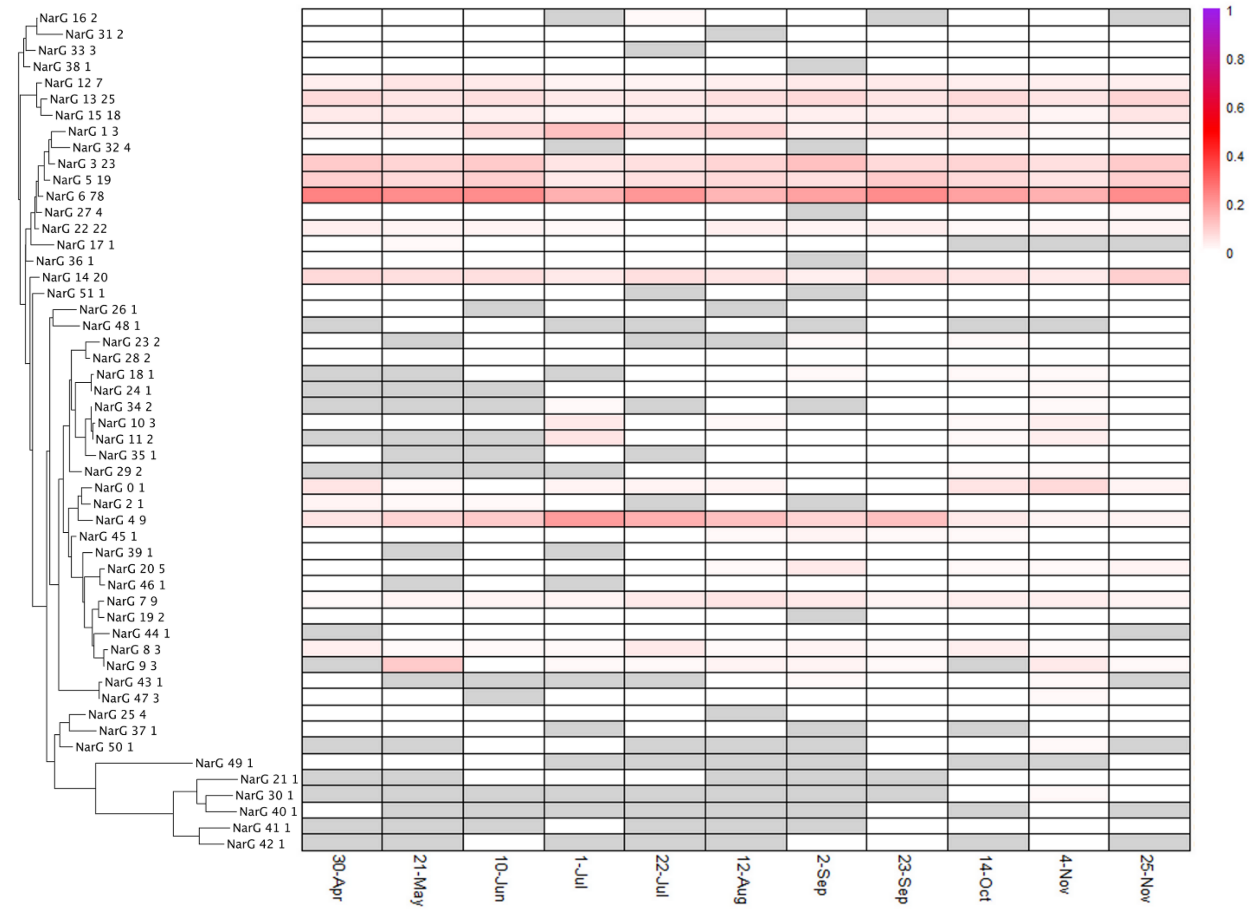


Fig S1A NarG gene phylotype distributions. Heatmap indicates the relative contribution of each phylotype (clustered at 90% AAID) to the total count; phylogenetic tree to the left of the heatmap demonstrates the diversity of phylotypes present.

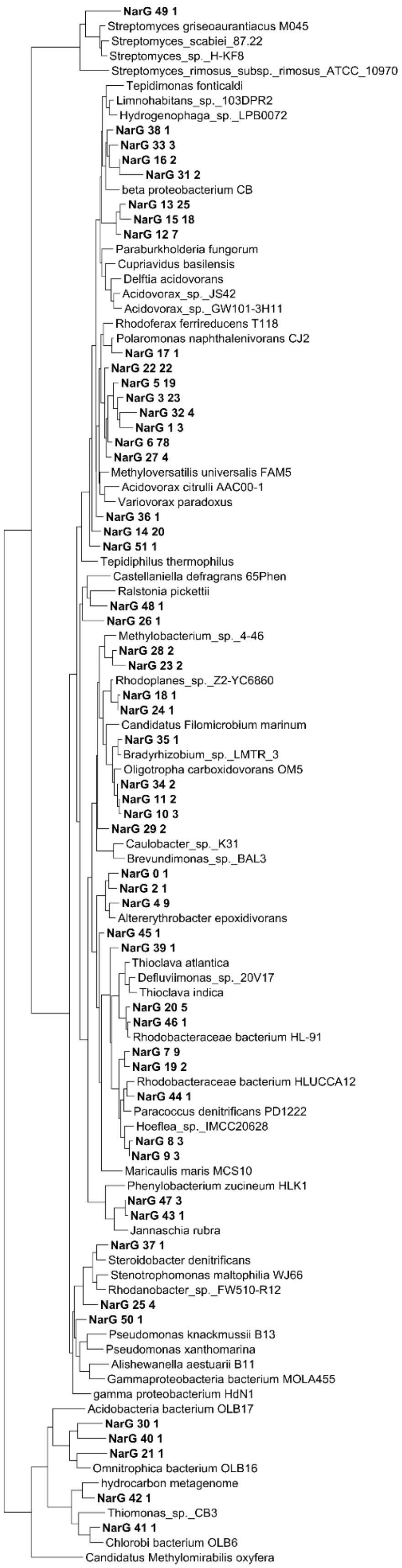


Fig S1B. NarG phylotype phylogenetic tree. Neighbor-joining tree of NarG Sequences assembled from the metagenomics samples and select NarG sequences from RefSeq.

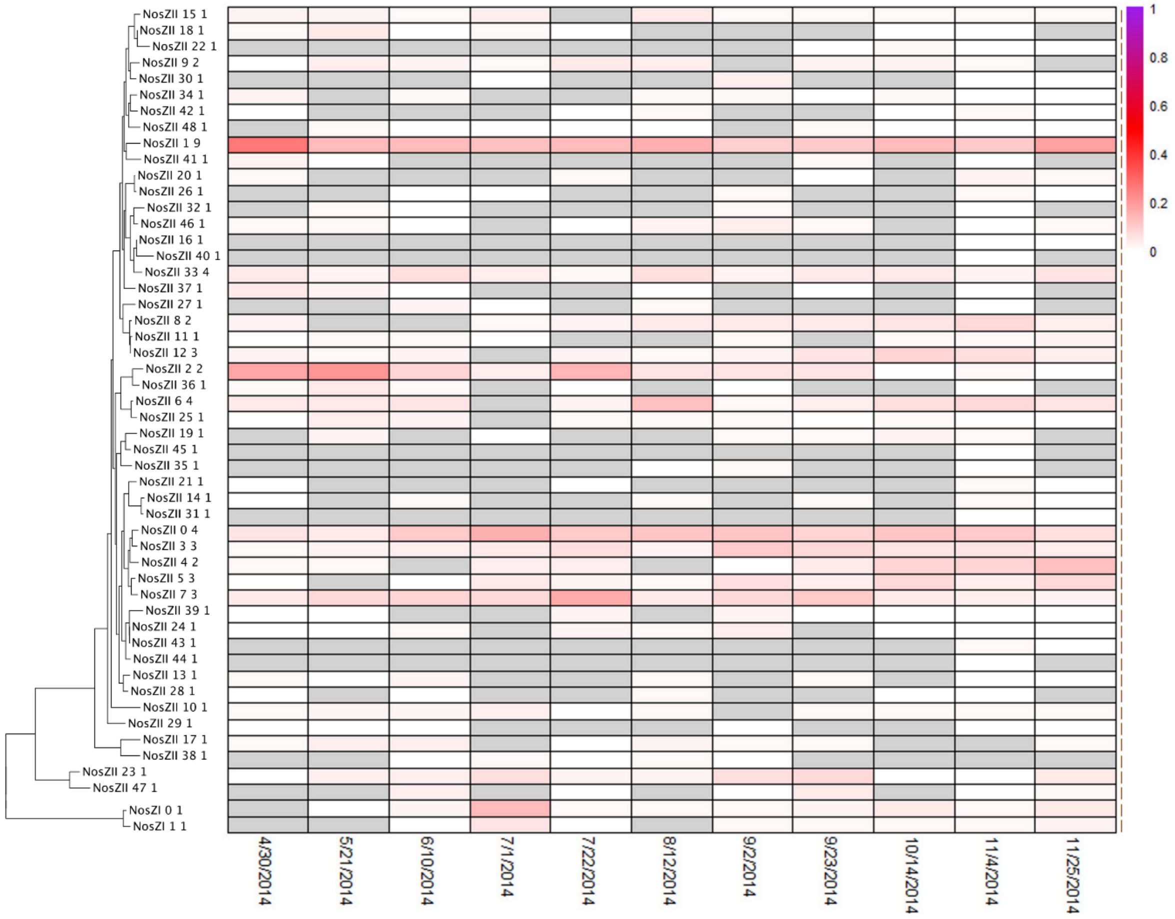


Fig S2A. nosZI and nosZII phylotype distributions. See Figure S1 for display details. Relative abundances for nosZI and nosZII were calculated separately.

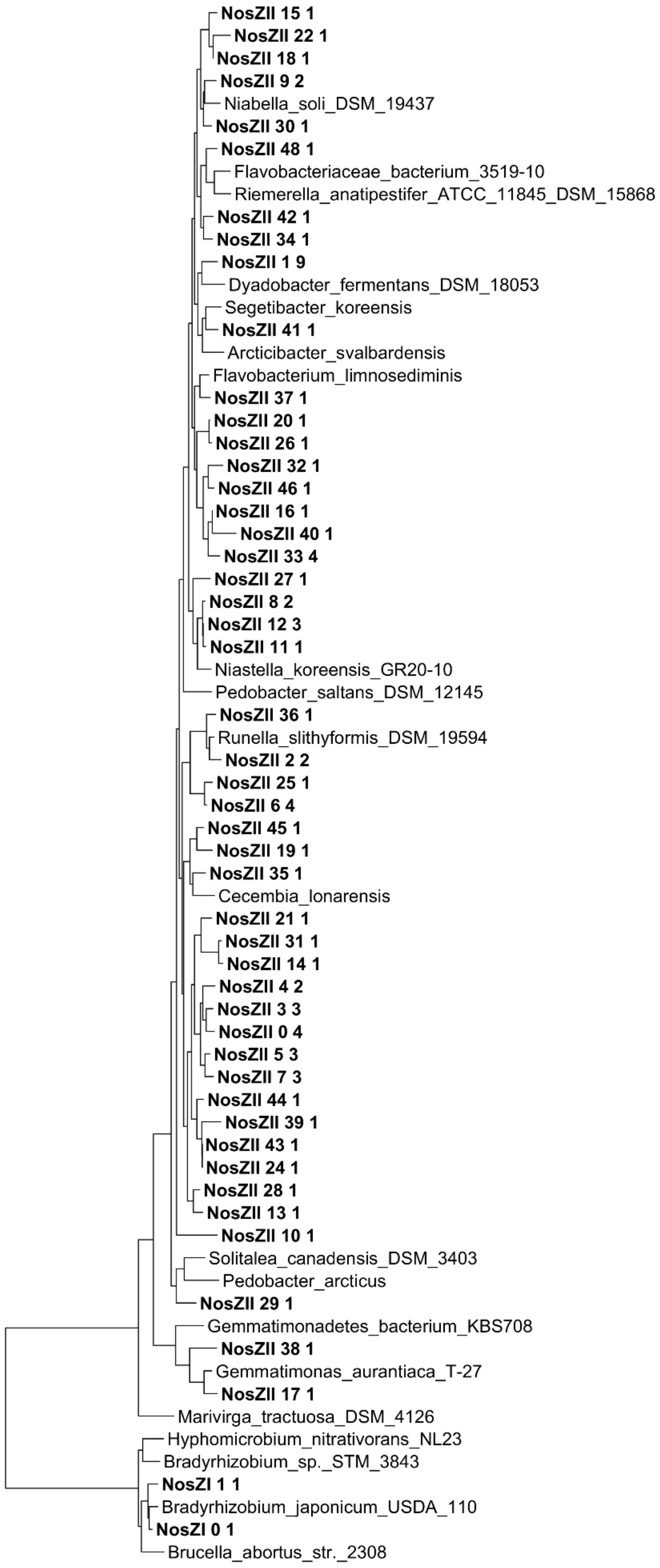


Figure S2B. NosZ phylogenetic tree. Neighbor-joining tree of NosZ Sequences assembled from the metagenomics samples and select NosZ sequences from RefSeq.

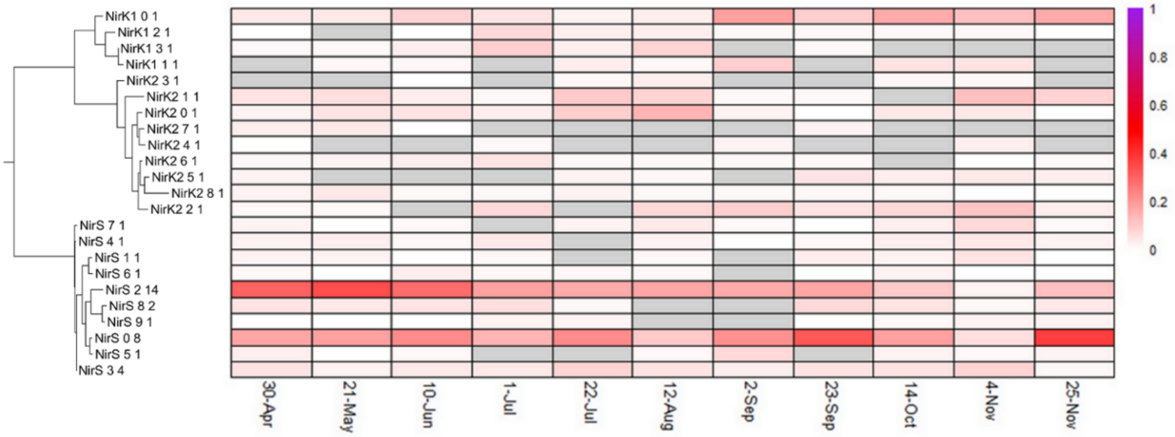


Fig S3A. NirK and NirS phylotype distributions. See Figure S1 for display details. Relative abundances were calculated grouping both gene types

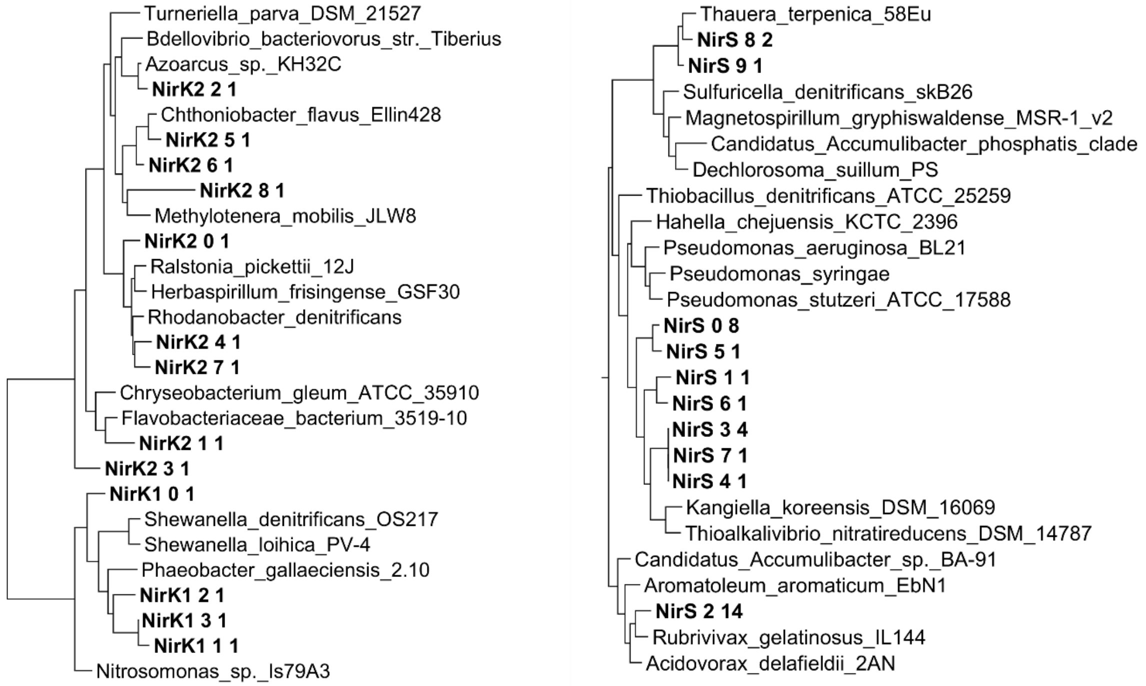


Fig S3B. NirK and NirS phylotype phylogenetic trees. Neighbor-joining tree of sequences assembled from the metagenomics samples and select sequences from RefSeq.

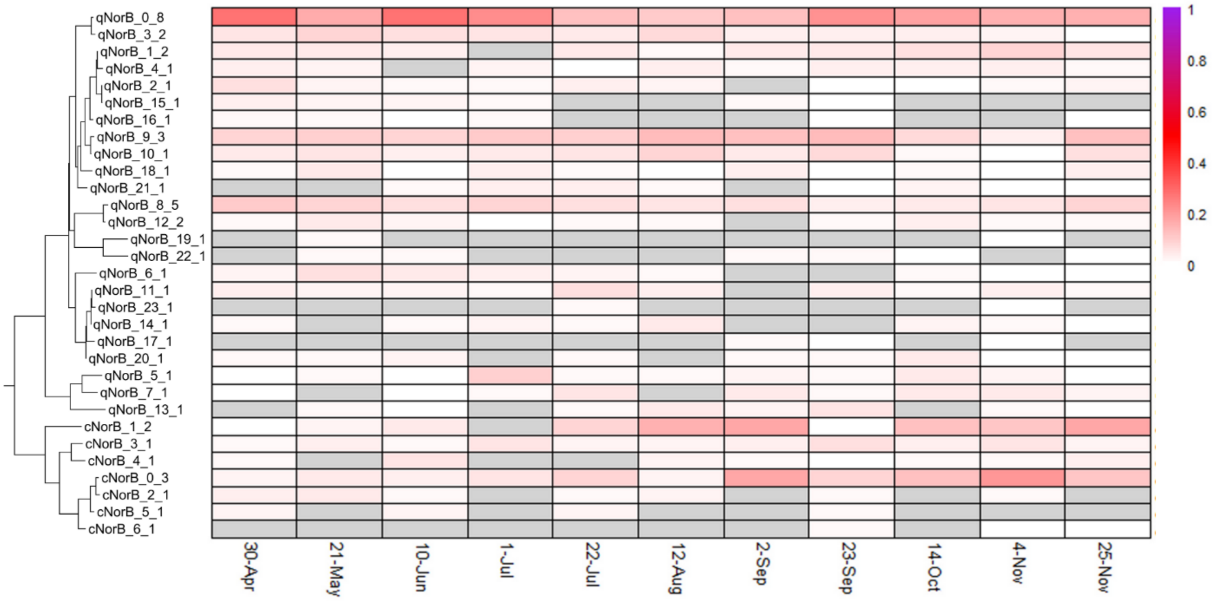


Fig S4A. NorB gene abundances and phylotype distributions. See Figure S1 for display details.

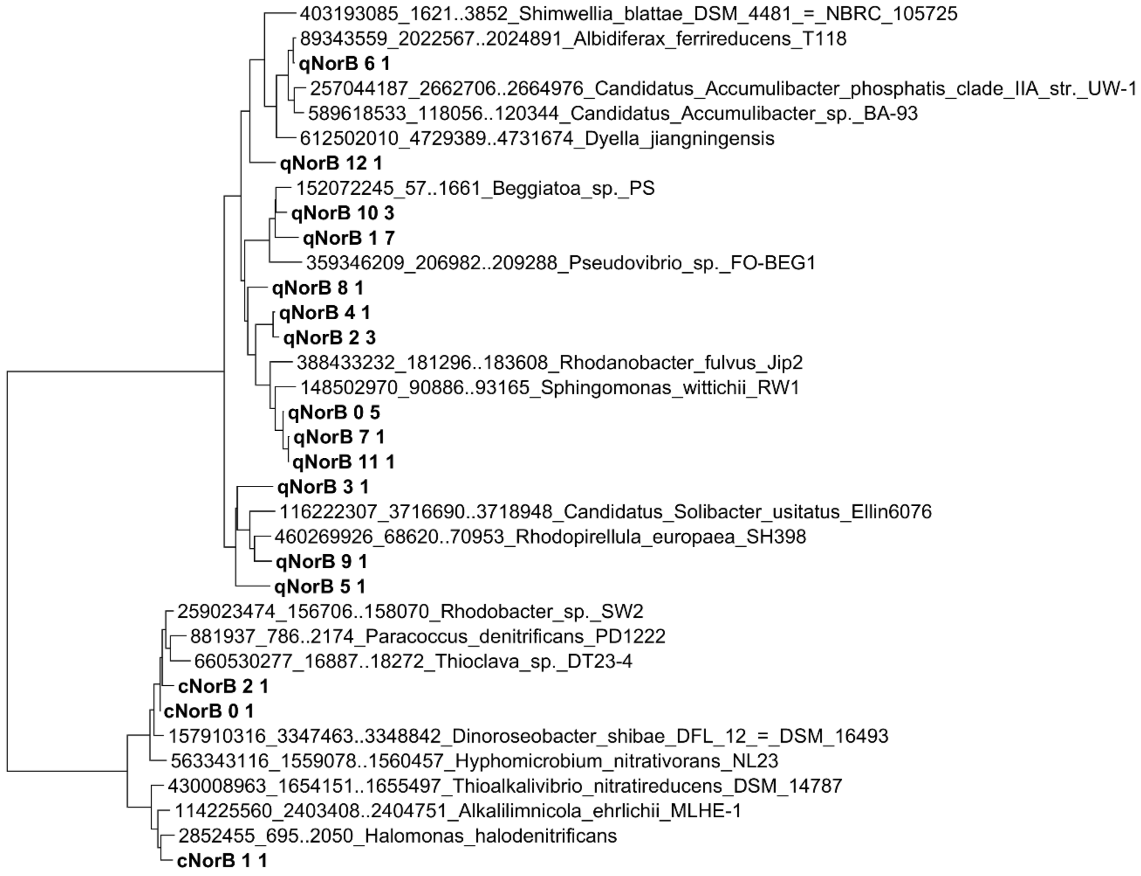


Fig S4B. NorB phylogenetic tree. Neighbor-joining tree of NorB sequences assembled from the metagenomics samples and select NorB sequences from RefSeq.

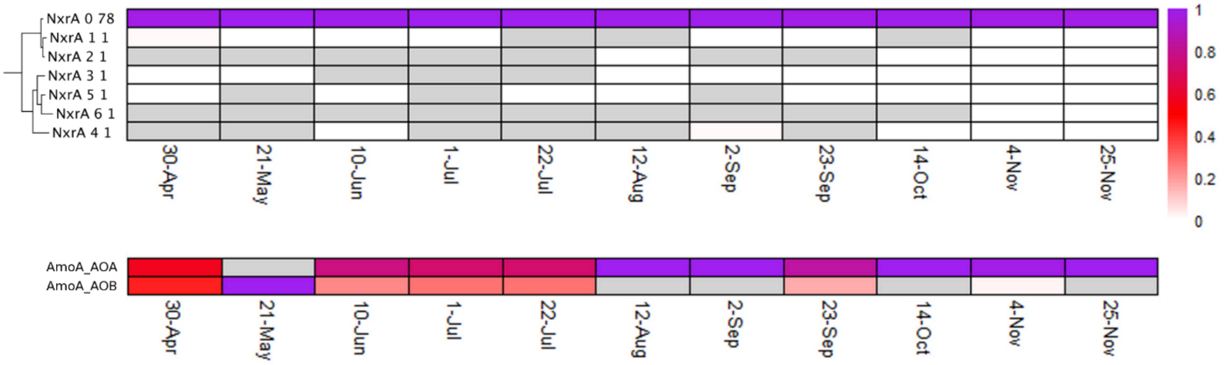


Fig S5A. NxrA and AmoA phylotype distributions. See Figure S1 for display details.

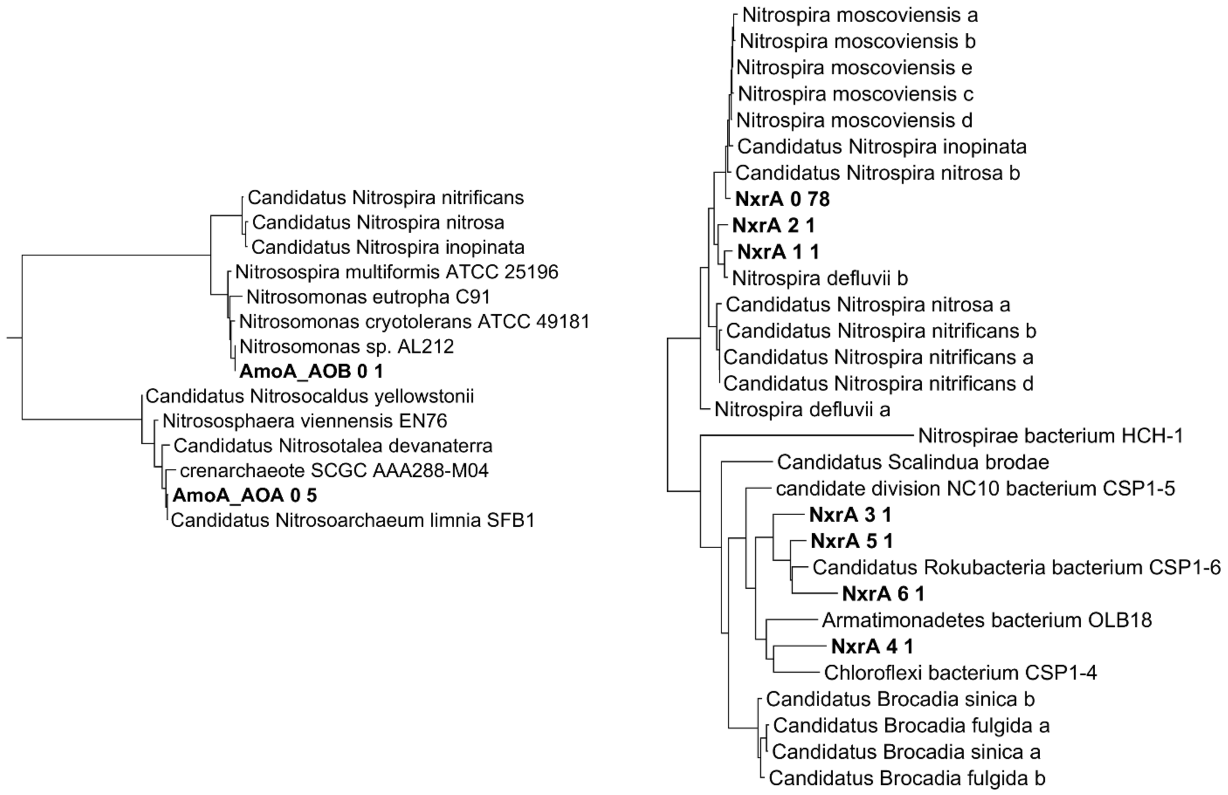


Fig S5B AmoA and NxrA phylogenetic trees. Neighbor-joining tree of sequences assembled from the metagenomics samples and select sequences from RefSeq.

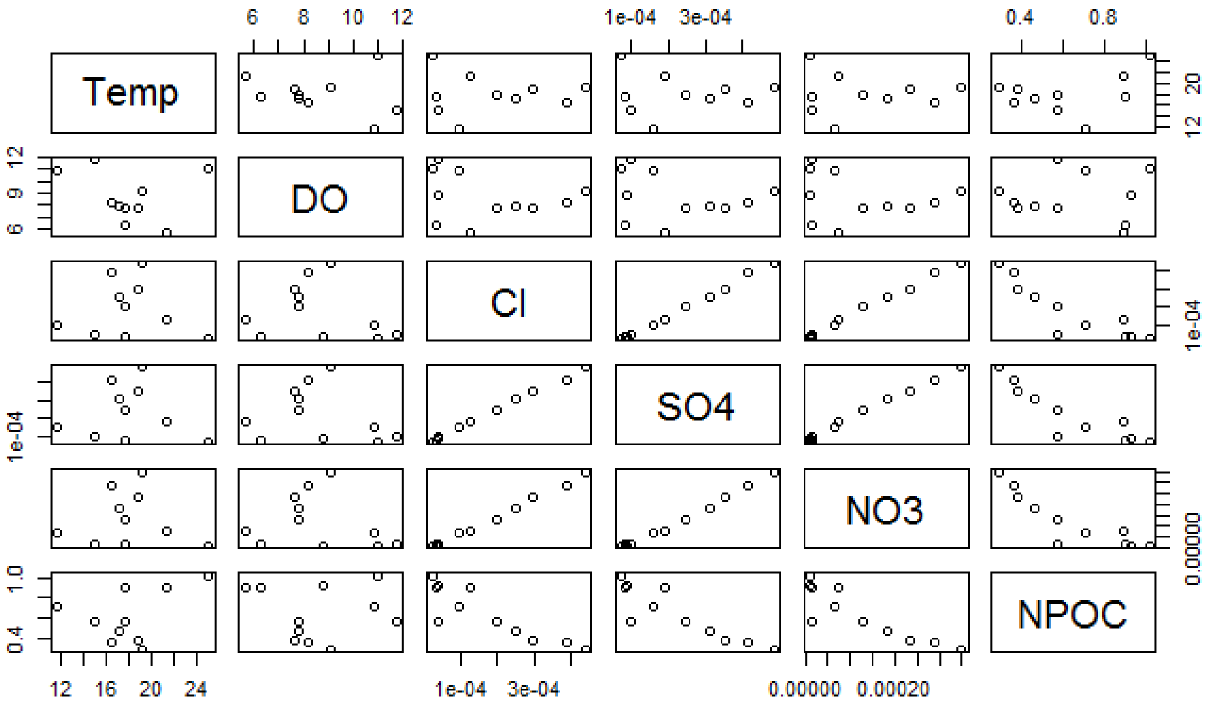


Figure S7. Environmental parameter correlation. Temperature (Temp), dissolved oxygen (DO), chloride ion concentration (Cl), sulfate concentration (SO4), nitrate concentration (NO3) and dissolved organic carbon (measured as non-purgeable organic carbon, NPOC) measurements were taken for all samples. Pair-wise correlation of observations were performed to determine the independence of the parameters.

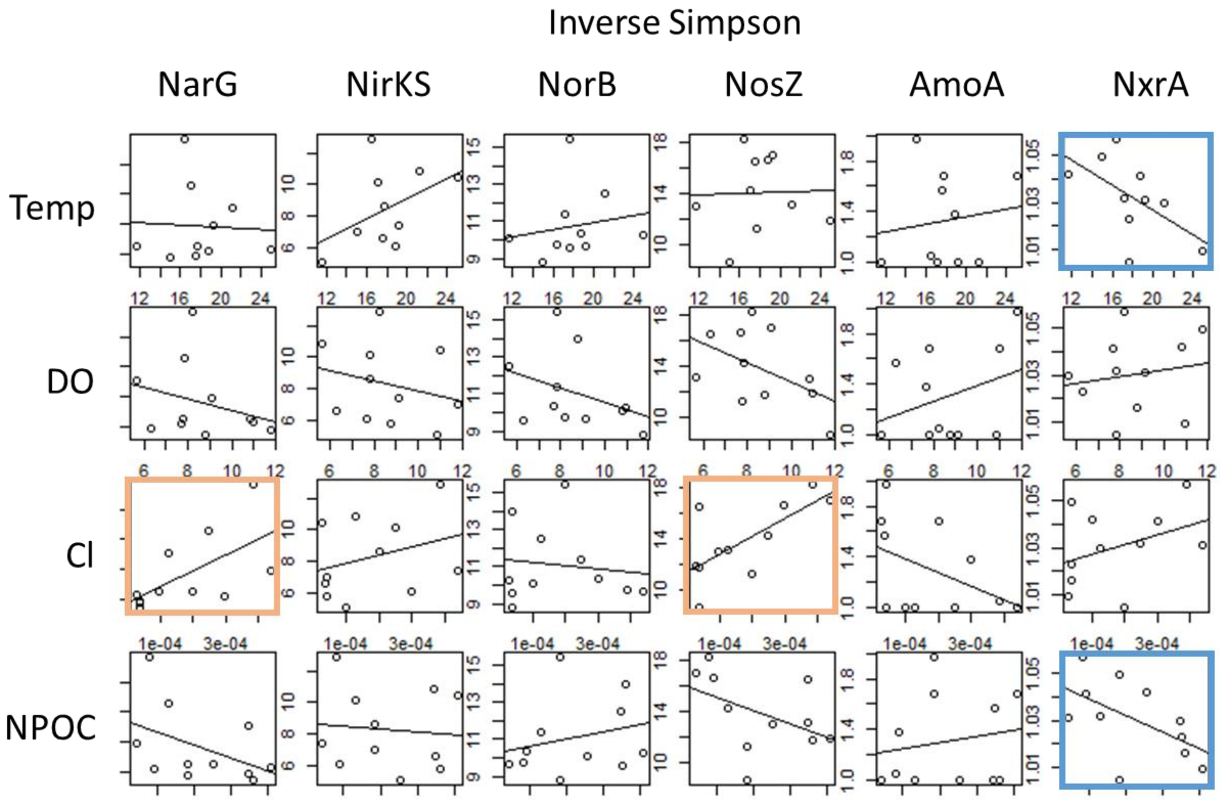


Figure S7. Environmental parameter vs diversity (as measured by the inverse Simpson statistic) linear regression analysis. Blue borders: $p < 0.10$; Orange borders: $p < 0.05$.

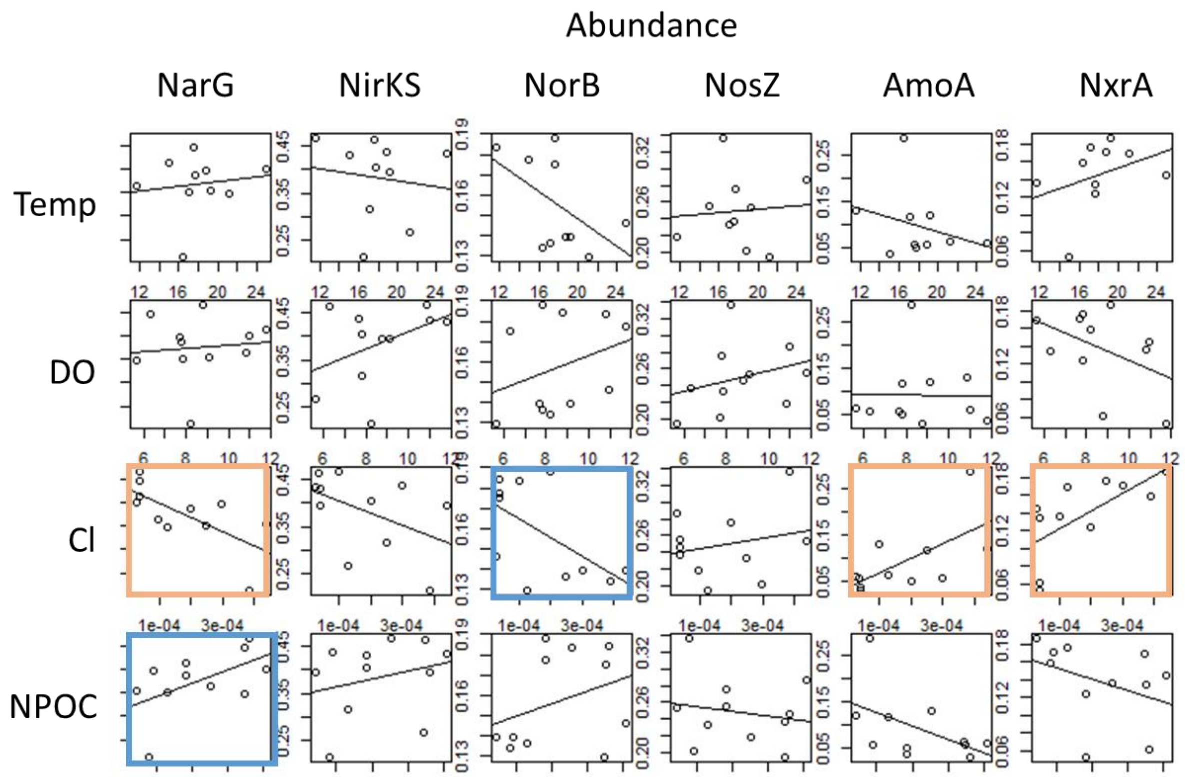


Figure S8. Environmental parameter vs gene abundance linear regression analysis. Blue borders: $p < 0.10$; Orange borders: $p < 0.05$.

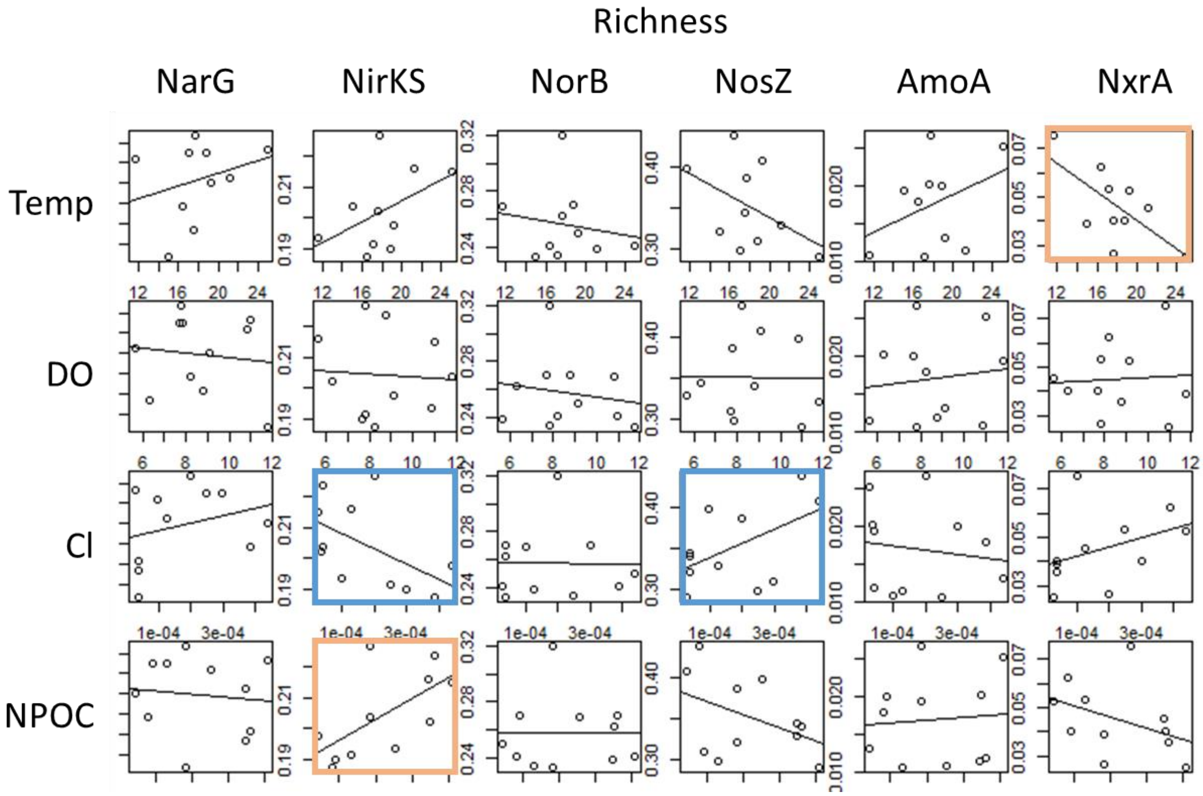


Figure S9. Environmental parameter vs richness linear regression analysis. Blue borders: $p < 0.10$; Orange borders: $p < 0.05$.
