## Supplemental Table S1 for "Temporal dynamics of nitrogen cycle gene diversity in a hyporheic microbiome"

**Table S1** Sequencing results.

| Piezometer | Date Collected | IMG project id | Total Raw Seqs |
|---|---|---|---|
| **T2** | 2014-04-30 | 1065985 | 48,020,244 |
| **T2** | 2014-09-23 | 1065999 | 30,613,936 |
| **T2** | 2014-11-25 | 1065975 | 16,734,972 |
| **T3** | 2014-04-30 | 1065997 | 33,660,036 |
| **T3** | 2014-09-23 | 1065989 | 19,489,174 |
| **T3** | 2014-11-25 | 1065993 | 18,218,980 |
| **T4** | 2014-04-30 | 1065983 | 48,412,732 |
| **T4** | 2014-05-21 | 1066005 | 25,681,518 |
| **T4** | 2014-06-10 | 1066001 | 33,554,732 |
| **T4** | 2014-07-01 | 1065991 | 19,761,996 |
| **T4** | 2014-07-22 | 1065977 | 16,159,946 |
| **T4** | 2014-08-12 | 1065987 | 19,728,928 |
| **T4** | 2014-09-02 | 1065979 | 16,770,554 |
| **T4** | 2014-09-23 | 1065995 | 36,917,050 |
| **T4** | 2014-10-14 | 1066003 | 33,326,776 |
| **T4** | 2014-11-04 | 1066007 | 62,894,882 |
| **T4** | 2014-11-25 | 1065981 | 60,562,476 |
